# A deep generative model of the SARS-CoV-2 spike protein predicts future variants

**DOI:** 10.1101/2023.01.17.524472

**Authors:** Rahul M. Dhodapkar

## Abstract

SARS-CoV-2 has demonstrated a robust ability to adapt in response to environmental pressures—increasing viral transmission and evading immune surveillance by mutating its molecular machinery. While viral sequencing has allowed for the early detection of emerging variants, methods to predict mutations before they occur remain limited. This work presents SpikeGPT2, a deep generative model based on ProtGPT2 and fine-tuned on SARS-CoV-2 spike (S) protein sequences deposited in the NIH Data Hub before May 2021. SpikeGPT2 achieved 88.8% next-residue prediction accuracy and successfully predicted amino acid substitutions found only in a held-out set of spike sequences deposited on or after May 2021, to which SpikeGPT2 was never exposed. When compared to several other methods, SpikeGPT2 achieved the best performance in predicting such future mutations. SpikeGPT2 also predicted several novel variants not present in the NIH SARS-CoV-2 Data Hub. A binding affinity analysis of all 54 generated substitutions identified 5 (N439A, N440G, K458T, L492I, and N501Y) as predicted to simultaneously increase S/ACE2 affinity, and decrease S/tixagevimab+cilgavimab affinity. Of these, N501Y has already been well-described to increase transmissibility of SARS-CoV-2. These findings indicate that SpikeGPT2 and other similar models may be employed to identify high-risk future variants before viral spread has occurred.

## 1 Introduction

The SARS-CoV-2 virus, causative agent of the COVID-19 pandemic, has claimed at least 5 and perhaps upwards of 18 million lives worldwide since it began circulating widely in early 2020 [1]. While successful vaccines have given clinicians and public health officials a powerful tool to curb viral spread and limit disease severity, SARS-CoV-2 has shown that it can adapt its molecular machinery through germline mutations that increase transmissibility and hamper immune surveillance [2, 3]. These mutated variants can lead to re-infection in those previously infected by the virus, as well as to breakthrough infections in vaccinated individuals. To prevent these infections, new vaccines have been developed to target the most prevalent and dangerous of these variants, and more are being developed as new variants continue to emerge [4].

The accurate and timely identification of variants that may give rise to new waves of infection remains an important component of the global effort to combating the evolving virus [5]. As part of this effort, several groups have compiled large volumes of viral sequencing data to track emerging variants. These sequences have been made easily accessible, for example, through the National Center for Biotechnology Information (NCBI) viral genomes resource, with unrestricted access worldwide [6]. To further improve preparedness for emerging variants, several computational tools have been developed to forecast the risk posed by an identified mutation, using a variety of techniques including combinations of epidemiological, biophysical, and/or deep learning approaches [7, 8, 9, 10, 11]. However, techniques to predict likely mutations *de novo*, before they have been observed in a human host, remain lacking.

This manuscript presents SpikeGPT2, a framework for the generation of SARS-CoV-2 spike protein sequences that can identify future variants of concern before they arise in the population. By applying transfer learning to a causal protein model (ProtGPT2), SpikeGPT2 can generate synthetic primary sequences for polypeptides of interest that vary meaningfully from reference data while adhering to the complex constraints governing protein structures [12, 13]. S protein receptor-binding motif (RBM) fragments synthesized by SpikeGPT2 contained mutations occuring with similar frequency and substitution distribution as sequences from the NIH Data Hub. Several S protein fragments synthesized by SpikeGPT2 contained point substitutions found only in held-out sequences, indicating that SpikeGPT2 was able to accurately predict mutations it had never seen before. A systematic comparison showed that SpikeGPT2 outperformed all other methods in minimizing the average number of predictions needed before accurately identifying a future mutation (measured by the number needed to simulate). Finally, the mutations predicted by SpikeGPT2 were analyzed using existing tools to identify potential variants of future concern.

## 2 Results

### 2.1 SpikeGPT2 model architecture and training

Similar to how large language models such as GPT2 can generate novel sequences of text and are trained on a next-word prediction task, large protein models such as ProtGPT2 can generate novel sequences of amino acids and are trained on a next residue prediction task (Figure 1A) [14]. To train the SpikeGPT2 model, a standard transfer learning approach was employed [15].

**Figure 1:**
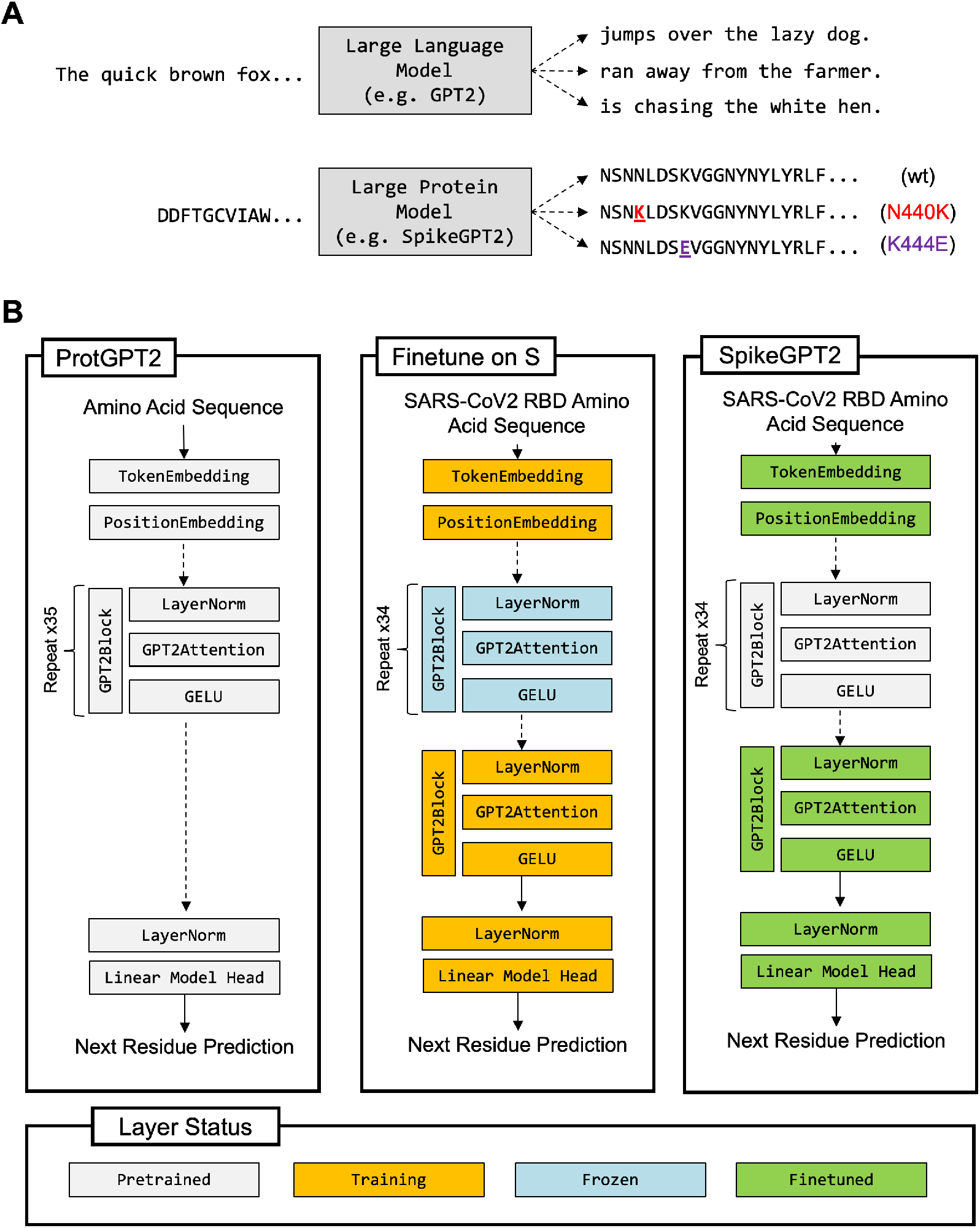
Model architecture and training strategy for SpikeGPT2. (A) Large protein models such as SpikeGPT2 can generate synthetic amino acid sequences given oligonucleotide prompts, analogous to large language models such as GPT2. (B) Fine tuning of a base model (in this case, ProtGPT2) on a specific task (in this case, SARS-CoV-2 S protein prediction) can be achieved by fixing (freezing) the majority of internal weights and training only a subset of layers in the network architecture. S: SARS-CoV-2 spike protein; wt: wild type.

SpikeGPT2 was derived from a base ProtGPT2 model pre-trained on a large corpus of primary amino acid sequence data that was subsequently fine-tuned on S protein sequences deposited in the NIH Data Hub before May 2021 (pre-5/21) (Figure 1B). After fine tuning, the model achieved 88.8% accuracy on a next-residue prediction task on a test set of S protein sequences deposited in the NIH Data Hub on or after May 2021 (post-5/21). As the model had never been directly presented with any amino acid sequences deposited after May 2021 during either pre-training or fine-tuning, any prediction of variants present only post-5/21 would represent true *de novo* variant prediction. In other words, from the perspective of the SpikeGPT2 model, all data in the post-5/21 test data set can be considered future data.

### 2.2 Generated sequences share properties with observed viral sequences

SpikeGPT2 was used to generate 10,000 primary amino acid sequences given a prompt of the 10 amino acids immediately preceding the RBM in the Wuhan-Hu-1 SARS-CoV-2 S protein reference (GenBank accession QHD43416.1) [16]. Amino acid point substitutions were identified after alignment of the synthesized fragments to the reference sequence. A total of 54 unique amino acid substitutions were identified in the generated sequences. A kernel density plot weighted by the frequency of point substitutions across locations in the SARS-CoV-2 RBM demonstrated several clear shared peaks between the pre-5/21, post-5/21, and SpikeGPT2-predicted (i.e. generated) data sets (Figure 2A). A correlation analysis of the frequency of variants shared between the pre-5/21, post-5/21, and predicted data sets also revealed a significant positive correlation between variant frequencies in each of these three data sets (Figure 3).

**Figure 2:**
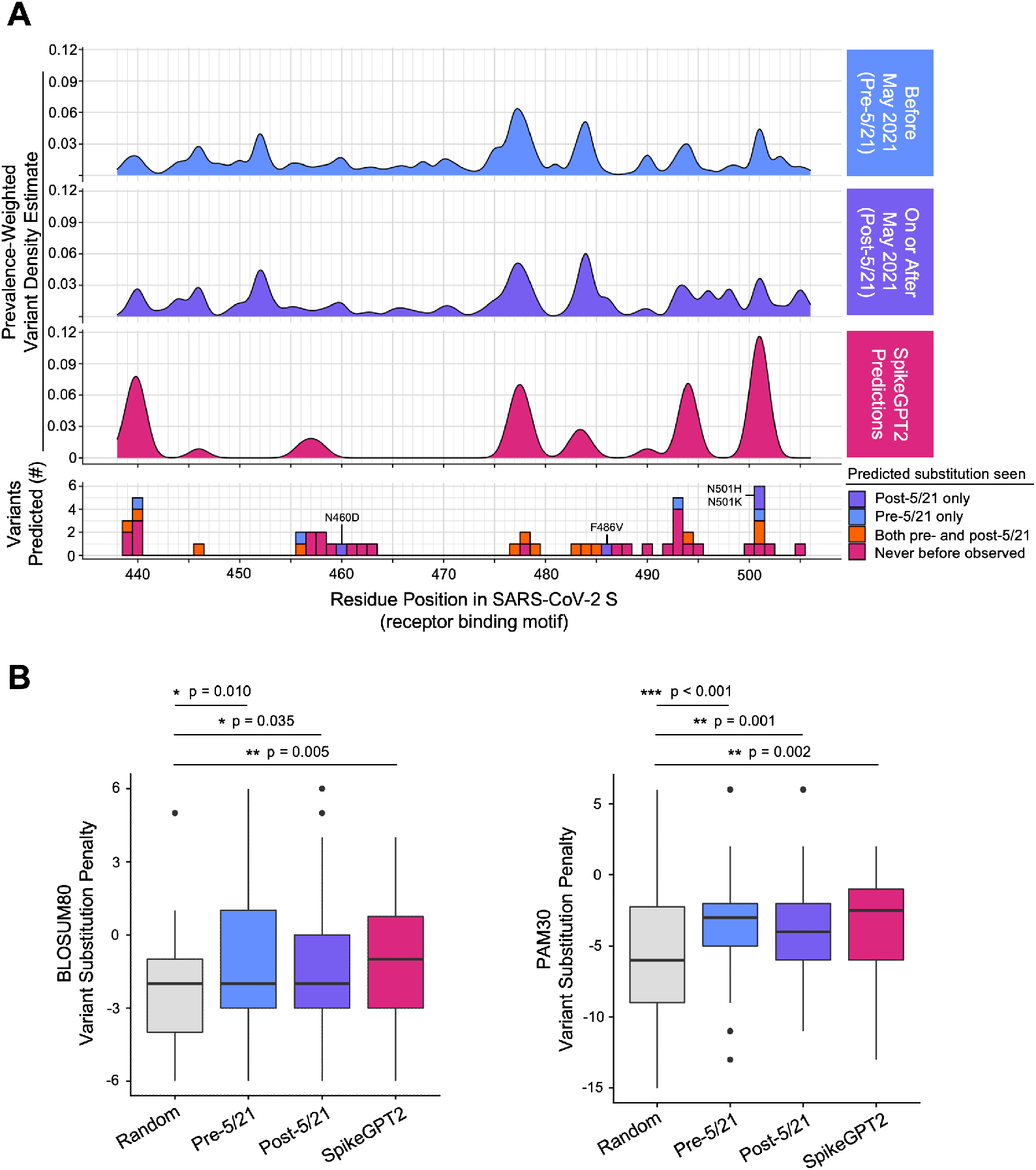
SpikeGPT2 generates RBM sequences that mirror the properties of circulating SARSCoV-2 RBM. (A) Gaussian kernel density estimate plots show that the observed population distribution of variants across positions in the SARS-CoV-2 S RBM both preand postMay 2021 are well approximated by SpikeGPT2 predictions. Specific variants generated by SpikeGPT2 present in the test set alone (valid future predictions) are labeled. (B) BLOSUM80 and PAM30 substitution matrices show that native SARS-CoV-2 S RBM and SpikeGPT2-predicted variants are less penalized than point substitution at random. S: spike protein; RBM: receptor binding motif.

**Figure 3:**
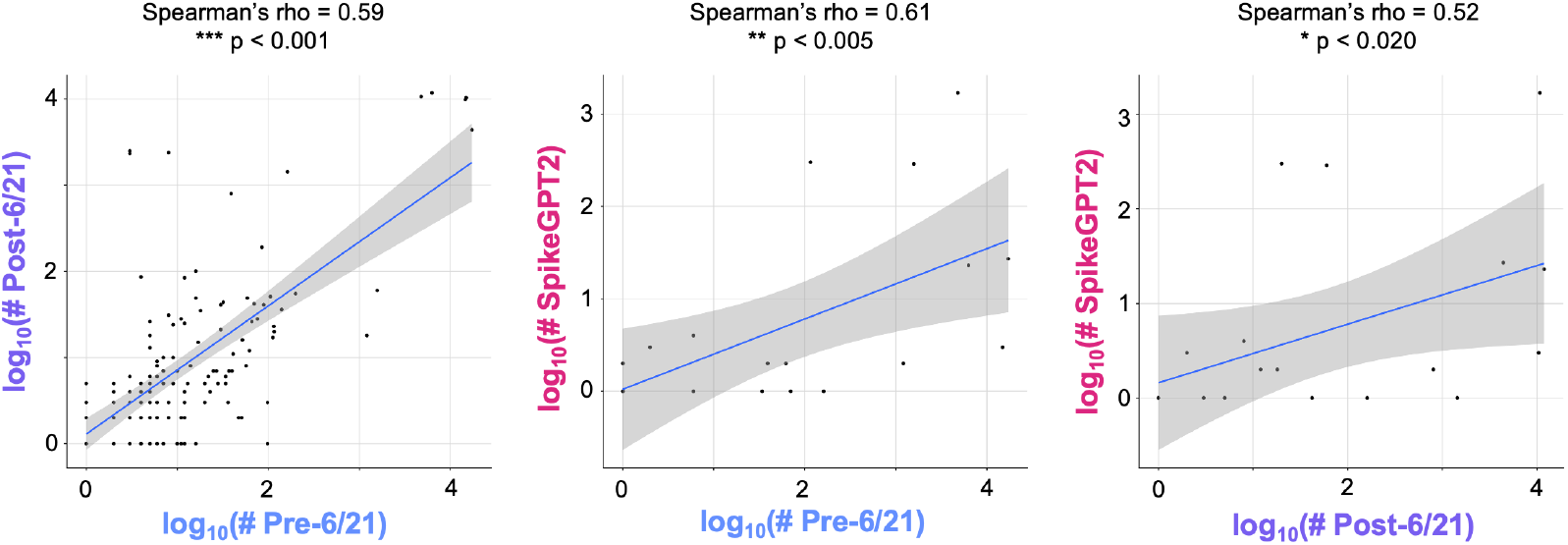
The population frequency of SARS-CoV-2 S RBM mutations is significantly correlated between all combinations of pre-May 2021 (pre-5/21), post-May 2021 (post 5/21) and SpikeGPT2predicted cohorts.

Standard substitution penalty matrices encoding the similarity of different amino acids were also used to assess the biological feasibility of point substitutions identified by the generative model. The total score of random point substitutions using both the BLOSUM80 and PAM30 matrices were significantly less than either the true point substitutions observed in the pre-5/21 (BLOSUM80 p = 0.010; PAM30 p *<* 0.001) and post-5/21 (BLOSUM80 p = 0.035; PAM30 p = 0.001) data sets, or the predicted point substitutions (BLOSUM80 p = 0.005; PAM30 p = 0.002) identified by SpikeGPT2 (Figure 2B).

### 2.3 SpikeGPT2 outperforms other methods of de novo variant prediction

To assess SpikeGPT2’s performance in *de novo* variant prediction, SpikeGPT2 was compared to three separate models in their ability to recover future variants in the SARS-CoV-2 S RBM. The comparator models were allowed access to the training (pre-5/21) data set to condition their predictions. For each model, a series of trials were conducted to measure the number of mutation simulations needed to recover a single mutation found in the test (post-5/21) data set. The number of trials needed was defined as the number needed to simulate (NNS), with a lower NNS indicating a more efficient mutation simulation process with higher fidelity to the underlying distribution of the mutation generating process.

The compared models were: (1) sample uniformly at random from the RBM locations, and return a point substitution at random (2) sample from the RBM locations, weighted by the known frequency of variants in the pre-5/21 dataset, and return a point substitution at random, and (3) sample from the RBM locations, weighted by the known frequency of variants in the pre-5/21 dataset, and return a point substitution weighted by the BLOSUM62 penalty vector for the germline amino acid. Each of the three models tested incorporated increasingly more prior knowledge into the variant simulation procedure. While the NNS decreased from Model 1 to Model 3, SpikeGPT2 achieved a NNS significantly lower than all other tested methods (Figure 4).

**Figure 4:**
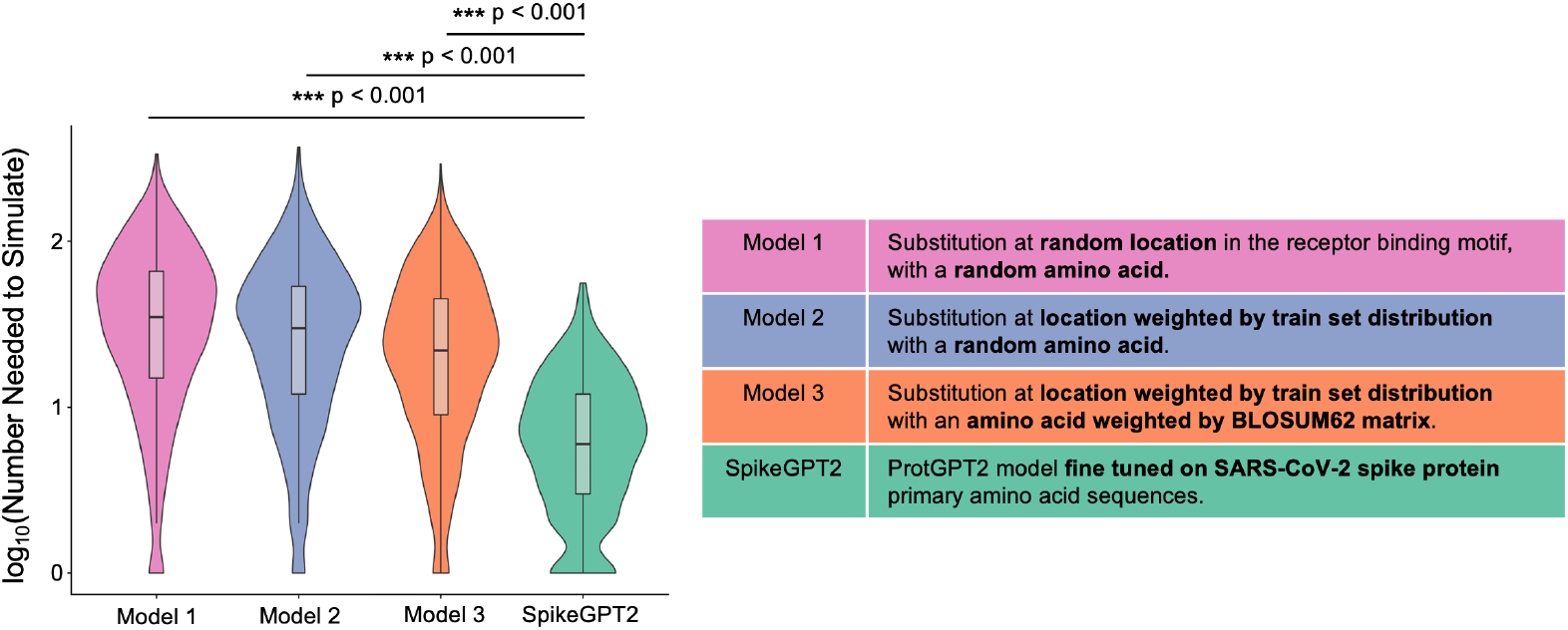
SpikeGPT2 outperforms all other benchmarked models in the efficient prediction of future SARS-CoV-2 S RBM substitutions. The number needed to simulate indicates the number of trials required to generate a single variant that was present only after May 2021 (post-5/21), given models with complete information prior to May 2021 (pre-5/21).

### 2.4 SpikeGPT2 predictions may be analyzed to identify potential variants of future concern

Once variants have been identified, either through synthetic or observational means, a number of existing tools and pipelines can be applied to assess the danger they may pose. Two means by which mutations can increase viral fitness are either through (1) increased transmissibility conferred by an increase in viral binding affinity for its host receptor (human ACE2 in the case of SARS-CoV-2), or (2) molecular evasion of immune or pharmacologic methods of curbing viral infection.

To assess these two axes of variation, mutation-induced binding affinity changes were simulated using MutaBind2 for the binding of SARS-CoV-2 S with tixagevimab and cilgavimab, therapeutic monoclonal antibodies used for pre-exposure prophylaxis (PrEP) against SARS-CoV2, as well as the binding of S to human ACE2 [17]. Mutations that both increased binding affinity for the S/ACE2 interaction, as well as decreased binding affinity for the S/tixagevimab+cilgavimab interaction were prioritized for further analysis (Figure 5A). Five such mutations were identified: N501Y, K458T, L492I, N439A, and N440G (Figure 5A-B). One of these mutations, N501Y, was present in both the preand post-5/21 datasets, and has been well described to increase SARS-CoV-2 S/ACE2 binding affinity [18, 19]. All five substitutions were close to the S/ACE2 interface, and visualizations were generated of the strain-minimizing rotamer at each mutated site using PyMOL.

**Figure 5:**
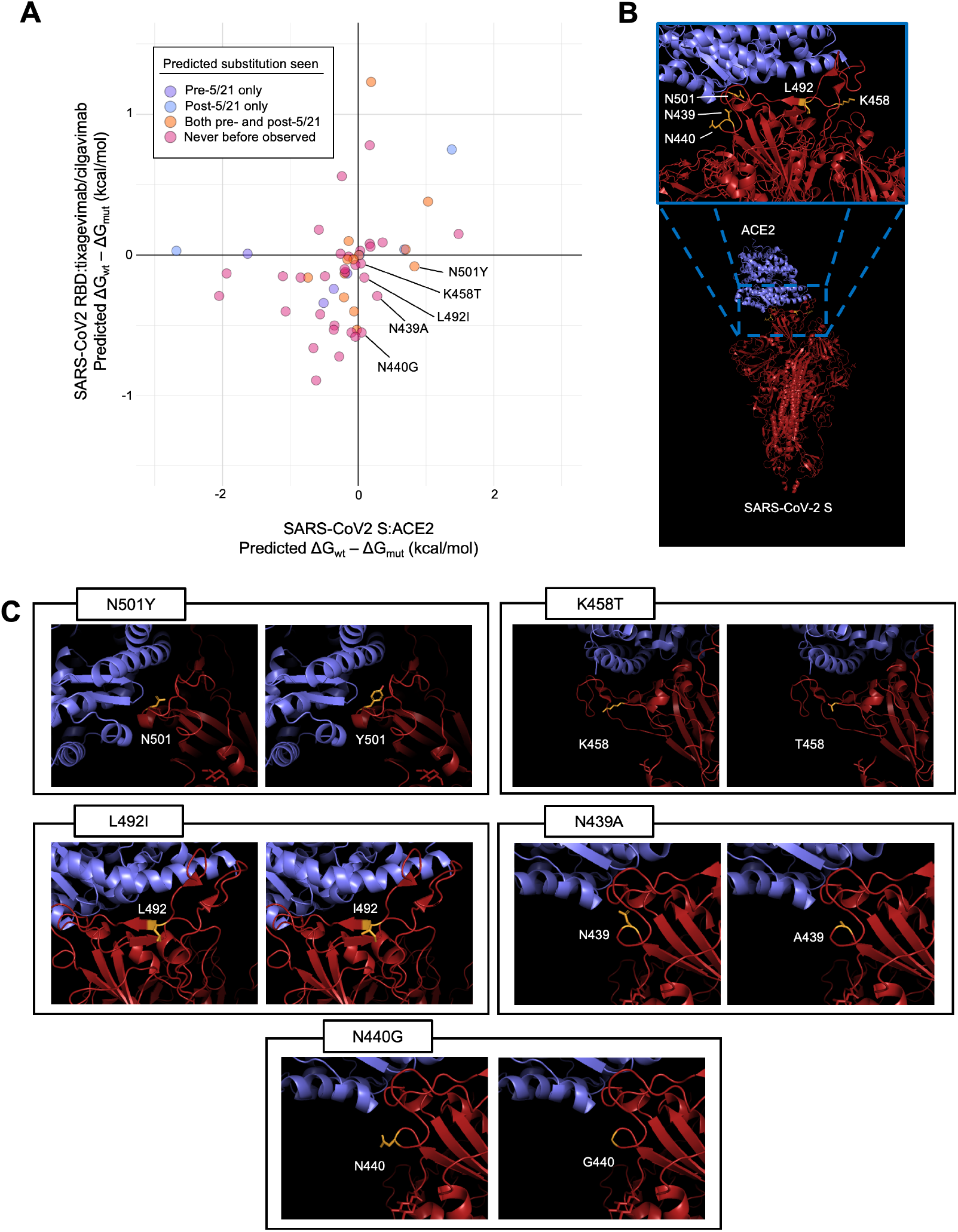
Analysis of predicted binding affinity changes induced by SpikeGPT2 identifies 5 mutations (N501Y, K458T, L492I, N439A, and N440G) that are predicted to both increase the binding affinity of SARS-CoV-2 S with human ACE2 as well as decrease the binding affinity of SARSCoV-2 S with the therapeutic antibodies tixagevimab and cilgavimab. (A) Scatter plot showing all SpikeGPT2-predicted mutations, and MutaBind2-generated binding energy changes. (B) Protein structure of SARS-CoV-2 S protein in complex with ACE2, showing the location of all mutations identified. (C) For each mutation, the predicted strain-minimizing rotamer is visualized.

## 3 Discussion

The identification of new mutations that may give rise to pathogenic lineages is essential in preparing for new waves of SARS-CoV-2, especially as the virus continues to circulate widely. While a number of *in silico* models have been developed to predict the risk of a given SARS-CoV-2 variant once it has been identified, approaches to accurately predict which variants are likely to develop in the future are limited, often involving direct scanning of the combinatorial space of possible primary sequences [20, 21]. Those methods which have explored new mutations have done so by heavily restricting the number of positions at which to consider new mutations, and/or by restricting the type of substitutions that may occur [22, 23]. Existing methods to scan the space of possible mutations suffer from poor efficiency: that is, they are able to predict new mutations that may eventually arise, but also generate a much larger pool of mutations that are unlikely to occur. SpikeGPT2 demonstrates significantly higher prediction efficiency compared to other models for exploring the space of possible variants, suggesting that it may substantially reduce the costs of investigating erroneous hits using laboratory methods.

While *in vitro* mutagenesis has been employed as an alternative to computational predictive approaches, there are several drawbacks to this strategy when dealing with infectious agents [24]. Experiments requiring the manipulation of functional viral particles are costly, requiring strict safety protocols and appropriate biohazard safety level (BSL) compliant facilities, that may not be easily available in low-resource settings [25]. Further, experiments that generate viable viral particles with increased pathogenicity run a theoretical risk of inadvertantly releasing engineered virions through breaches in protocols, causing laboratory-acquired infections that may spread to the surrounding communities [26]. Computational approaches to mutagenesis such as SpikeGPT2 are cheaper and safer than biological approaches for initial exploratory evaluations, and computational predictions can be validated through targeted *in vitro* investigations.

Large language models such as GPT2, upon which SpikeGPT2 is based, have proven themselves to be incredibly versatile, with the same model backbone capable of performing such varied tasks as code generation, question answering, and sentiment analysis, given the appropriate prompts. Likewise, while SpikeGPT2 was tested in this manuscript against a limited benchmark of detecting point substitutions in the SARS-CoV-2 S RBM, the model itself is not limited to these tasks. Indeed, the synthetic peptide fragments generated by SpikeGPT2 may be used to simulate single or multiresidue insertions, deletions, and other mutation patterns. While the SpikeGPT2 model was given a prompt of the last 10 amino acids immediately preceding the RBM, simple modification of the prompt can also allow it to explore other S protein subdomains. For example, by providing a blank prompt, SpikeGPT2 will begin synthesis at the N terminal domain (NTD), another important locus of S protein variation playing a key role in epitope recognition [27].

Though SpikeGPT2 shows promise as a method to generate new viral sequences, several important limitations to the method remain. The generative model is non-deterministic and may require many rounds of simulation to generate predicted mutations. In this study, 10,000 synthetic RBM fragments were generated, resulting in only 54 single residue substitutions. Additionally, generative methods are known to be highly sensitive to the choice of prompt used to initiate synthesis. These phenomena have been studied extensively in large language models trained on natural language corpora by the growing field of prompt engineering [28]. Finally, though there has been major progress in adding explainability to deep learning models through the use of attention and other approaches, the reasoning behind specific predictions generated by large neural networks such as SpikeGPT2 remains difficult to interpret [29].

In summary, this study demonstrates that fine-tuning of large generative models can be used to successfully predict future SARS-CoV-2 S RBM mutations. This investigation also lays the groundwork (and provides a code base) for using similar fine-tuning approaches to predict variability within other important proteins in human pathogens, viral or otherwise. By anticipating important variants before they occur, therapies may be designed to be more robust to future pathogenic adaptations, which may improve outcomes in the treatment of many infectious diseases.

## 4 Methods

### 4.1 Data Preparation

All surface glycoprotein primary sequence data from the NIH SARS-CoV-2 Data Hub was downloaded in FASTA format. The data were segmented into two sets: a training set containing all sequences collected before May 2021 (pre-5/21), and a training set containing all sequences collected afterward (post-5/21). As the loaded ProtGPT2 model had been pretrained on the UniRef50 (version 2021 04) dataset, this ensured that there was no possibility that variants identified after May 2021 could have been directly shown to the model during either the pretraining or the fine-tuning procedures. Due to resource constraints, subsampling was performed: from the full test and train data sets, 20,000 training samples and 10,000 test samples were chosen uniformly at random for model fine-tuning.

### 4.2 Model Training

The ProtGPT2 model from the HuggingFace transformers model hub was loaded using the `transformers` package from the python package index (PyPI) [30, 13]. Weights for all but the last GPT2 transformer layer were frozen during the fine-tuning process (Figure 1B). Fine-tuning was performed for 3 epochs through the training set, using a single NVIDIA A100 GPU. Hyperparameters were set as follows: learning rate of 10^−5^, training and evaluation batch sizes of 16, random seed of 42, ADAM optimizer with *β*_1_ = 0.9, *β*_2_ = 0.999, and *ϵ* = 10^−8^, and a linear learning rate scheduler. Hyperparameter optimization was performed using manual exploration of the learning rate and batch size parameters. Other hyperparameters were not modified from default values as per the HuggingFace transformers recommended settings. Different subsamples of the training and test data were used during hyperparameter optimization.

### 4.3 Synthetic Receptor-Binding Motif Generation

After model fine-tuning and evalutation on the test set, the model was used to generate 10,000 amino acid sequences with the prompt “`DDFTGCVIAW`”: the last 10 amino acids immediately preceding the receptor-binding motif (RBM) of the spike glycoprotein spanning from position 438-506 [5, 31]. No other prompt configurations were tested.

### 4.4 Point Substitution Calling

All FASTA sequences in the full (not subsampled) train (pre-5/21) and test (post-5/21) data sets as previously described were aligned to the Wuhan-Hu-1 SARS-CoV-2 primary amino acid sequence for S protein using the BioPython `Align` utilities [16, 32]. To avoid multiple alignments when aligning RBM-proximate fragments to the full surface glycoprotein primary amino acid sequence, an open gap score of -2 was imposed, and the BLOSUM62 matrix was employed for alignment [33]. All ungapped alignments were examined and point substitutions were called by direct string comparison. An analogous procedure to call point substitutions in the synthetic RBM fragments from SpikeGPT2 was also performed.

### 4.5 Substitution Frequency Analysis

The population frequencies of mutations in the full train (pre-5/21), test (post-5/21) and SpikeGPT2generated data sets were calculated and frequency-weighted Gaussian kernel density estimate plots were generated for each cohort to visually assess regions along the SARS-CoV-2 RBM that had the highest degree of variability. These kernel density estimate plots were visually inspected to assess for shared peaks between the distributions in the three data sets.

To formally assess the concordance between the population frequency of different variants, pairwise correlation analysis was performed. For each possible pairing of the train (pre-5/21), test (post-5/21) and SpikeGPT2-generated data sets, mutations found in both datasets were selected and Spearman correlation was calculated. The p-value of the correlation was also reported. Mutation frequencies were visualized using a 2D scatter plot, and a linear model was fit to each data set and plotted along the same axes to highlight the main relationships within the data.

### 4.6 Substitution Feasibility Analysis

To assess the validity of the point substitutions predicted by SpikeGPT2, the penalty scores of all substitutions were calculated using both the BLOSUM80 and PAM30 amino acid similarity matrices [33, 34]. A set of random point substitutions equal in number to the predicted substitutions from the ProtGPT2 model was also generated as a comparison group. For both the BLOSUM and PAM matrices, a less acceptable substitution is indicated by a more negative value. The BLOSUM80 and PAM30 distributions generated between the random point substitutions, and the train, test, and SpikeGPT2-predicted data sets were compared using two-tailed Wilcoxon rank sum tests.

### 4.7 Comparison of SpikeGPT2 to other mutation-generating models

The predictive performance of the generative model was assessed against three other methods of predicting new mutations *de novo*: (1) uniformly random distribution of variants across the simulated region, (2) test set frequency weighted distribution of variants across the simulated region with a random amino acid substitution change, and (3) test set frequency weighted distribution of variants across the simulated region with an amino acid change weighted by the row corresponding to the reference amino acid in the BLOSUM62 matrix. As the BLOSUM62 matrix contains negative entries, a transformation procedure was applied to each row of the BLOSUM62 matrix for Model 3: the minimum value of each row was subtracted from the elements of the corresponding row of the matrix, and the matrix row-sums were normalized to 1 (to indicate sampling probabilities).

For each of the above procedures, mutations were simulated until a mutation that was present in the test set (post-5/21) but not in the train set (pre-5/21) was encountered. The number of simulations required to reach such a mutation was recorded as the number needed to simulate (NNS). Simulations yielding a mutation that was present in the train set (pre-5/21) were discarded and not counted towards the NNS score. For each model, 999 trials were performed and the distribution of NNS values obtained were plotted. For SpikeGPT2, bootstrapped sampling of the predicted mutations was used to generate 999 trials of the NNS score. A two-tailed Welch’s t-test for independent samples was used on the log-transformed NNS distributions to assess for significant location shift across all pairs of models. Bonferroni correction was applied for multiple comparisons.

### 4.8 Binding Energy Analysis

The changes to binding energy generated by the predicted point substitutions between the SARSCoV-2 S glycoprotein and other binding targets of interest was assessed using MutaBind2 [9]. MutaBind2 enables the direct estimation of the change in binding free energy (ΔG) given a pre-defined protein structure and a given amino acid substitution. To assess the binding between SARS-CoV-2 spike and the ACE2 receptor, a cryo-EM solved protein structure of the S-ACE2 complex available on the RCSB Protein Data Bank at accession 7DF4 was used [35]. To assess the potential for mutations in SARS-CoV-2 spike protein to escape from existing therapies, a cryo-EM solved protein structure of SARS-CoV-2 Spike RBD in complex with tixagevimab/cilgavimab at accession 8D8Q was used [36]. Point substitutions that caused both a predicted increase in binding affinity between S and ACE2 as well as a predicted decrease in binding affinity between S and tixagevimab/cilgavimab were highlighted as potentially high-risk variants. These selected variants were plotted using PyMOL and strain-minimizing mutant amino acid rotamers were determined using the PyMOL mutagenesis tools.

### 4.9 Statistics and Visualizations

Plots were generated using the `seaborn` and `matplotlib` libraries for python as well as the `ggplot2` library for R [37, 38, 39]. Molecular visualizations were performed using the opensource version of PyMOL. Selected variants were visualized in their predicted strain-minimizing rotamer configuration using the mutagenesis wizard in the base PyMOL installation. All statistical testing was performed using the base language utilities in R. Specific tests employed are detailed in the relevant sections of the preceding methods.

## Data availability statement

All data used in the manuscript are publicly available. SARS-CoV-2 S protein amino acid sequences arew available from the NIH SARS-CoV-2 Data Hub [6]. All protein structure data is publicly accessible under the protein data bank (PDB) at accessions 7DF4 and 8D8Q [40].

## Code availability statement

Code is available in a public GitHub repository under the Creative Commons AttributionNoncommercial-ShareAlike 4.0 License at:

https://github.com/rahuldhodapkar/PredictSARSVariants

A full copy of the fine-tuned model is made available for use at:

https://huggingface.co/rahuldhodapkar/protgpt2-finetuned-sarscov2-rbd

